# Caspase-3/Drice as a critical regulator of actin dynamics through its dual control of small RhoGTPase family and Gelsolin in the Malpighian tubules of *Drosophila*

**DOI:** 10.1101/2025.10.01.679759

**Authors:** Saurabh Chand Sagar, Madhu G Tapadia

## Abstract

Caspases are executioner enzymes primarily known for their role in driving programmed cell death. However, they are now also known to regulate plethora of non-apoptotic functions such as proliferation, differentiation, endocytic trafficking, cell polarity, morphogenesis, inflammatory response and immune response. Here, in this study we report the role of *Drosophila* caspase 3, Drice, in spatial and dynamic regulation of actin filaments during development and proper functioning of the Malpighian tubules (MTs) of *Drosophila melanogaster*. Previously, we showed that Drice is crucial for the morphogenesis of the MTs, and its absence results in erroneous RhoGTPase signaling driving disarray in actin organization, this led to the formation of multiple fluid filled cysts in tubules. In the present study, we show that altered expression of two member of the Rho family of small GTPases, Rho1 and CDC42, perturbs the downstream signaling cascade. Reduced expression of Rok aborts the Rho1 signaling in Drice null mutants, whereas enhanced expression of CDC42 results in Arp2/3-driven hyper-polymerization of actin filaments. We compared Rho-GTPase interacting partners in *Oregon R+* (Control) and Drice mutants, and identified the absence of Gelsolin-Rho1 interaction in Drice mutants. Moreover, its expression was significantly downregulated, in the MTs. These factors collectively influence the F-actin to G-actin ratio, which significantly affects actin organization in the MTs. These results demonstrate the crucial role of caspase-3/Drice in maintaining actin homeostasis and tubule morphogenesis. This study highlights the importance of caspase activity beyond apoptosis and its potential function in the developmental and physiological processes.

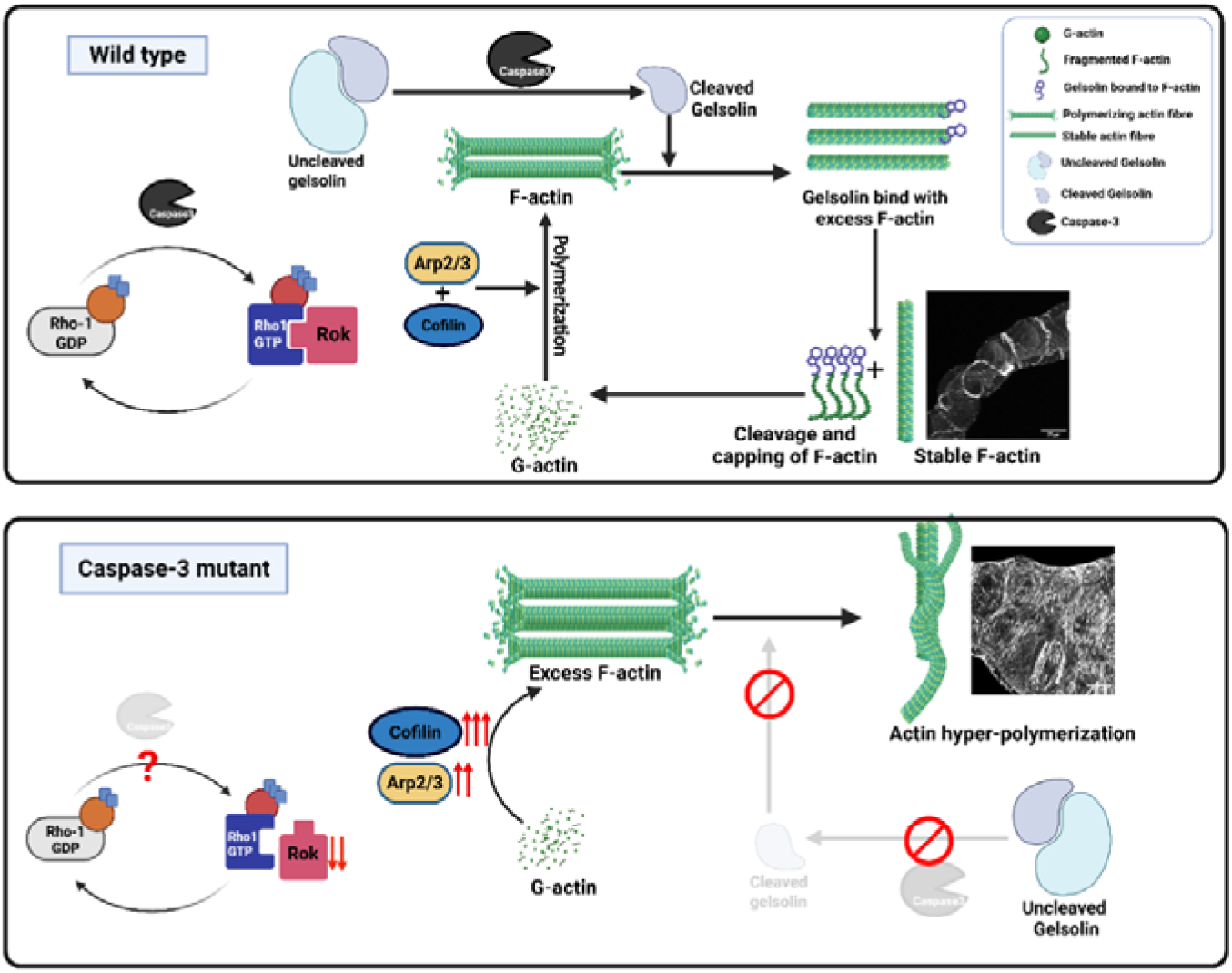

## Introduction

Morphogenesis is a fundamental process in development, as it defines how cells acquire their shape and spatial organization—both of which are indispensable for conventional morphology and organ physiology. This transformation from an undifferentiated cell mass into specialized functional structures relies on the coordinated remodeling of individual cell shapes and positions (1, 2). A critical element of this remodeling is the close interplay between the plasma membrane and the actin cytoskeleton, which together safeguard cellular integrity while enabling morphological changes and cellular movement/motility (3, 4).

Actin filaments provide the structural framework, and its dynamic behavior of switching between monomeric (G-actin) and filamentous (F-actin) states, support cell migration, shape transitions, tissue metamorphosis and differentiation (5, 6). Typically, cytosolic G-actin levels are high, but in response to signaling cues it polymerizes into F-actin; conversely, F-actin can depolymerize when required by the cell (7, 8). This dynamic equilibrium is regulated by a diverse set of actin-binding proteins (ABPs), including filament nucleators (such as formins and spire proteins), branching regulators (e.g., Arp2/3 activators), severing factors (such as gelsolin and villin), and proteins that cap or stabilize filaments (9–13). The recruitment and activity of ABPs are directed by upstream signaling networks, such as PI3K-AKT pathway, Rho family proteins, Yorkie-Hippo pathway, MAPK pathway, mTOR, Calcium dependent regulation, Wnt signaling etc. Of these, RhoGTPase family represent a central hub in actin regulation (14, 15). These molecular switches alternate between active GTP-bound and inactive GDP-bound states, under the control of guanine nucleotide exchange factors (GEFs), GTPase-activating proteins (GAPs), and RhoGDIs (16, 17). Among the more than 20 known Rho GTPases, RhoA, Rac1, and CDC42 are particularly important in actin regulation (18, 19). RhoA stimulates stress fiber assembly by activating effector proteins that promote filament bundling (20). In contrast, Rac1 drives lamellipodia formation, and CDC42 triggers filopodia extension, largely through Arp2/3-mediated actin branching and elongation (21). Beyond their individual roles, RhoA, Rac1, and CDC42 engage in reciprocal regulation, creating a dynamic balance that ensures robust yet flexible control of cytoskeletal architecture and morphogenetic progression.

Caspases, activator and executioner, are best known for their key roles in programmed cell death (22). However, emerging evidence indicates non-apoptotic functions for certain caspases, including caspase-11, which has been implicated in cell migration and actin depolymerization (23, 24). Several studies highlight functional interactions of caspases with signaling molecules that regulate cytoskeletal dynamics directly or indirectly. For example, caspase-3 indirectly influences actin organization by modulating Rho-associated kinase (Rok), a downstream effector of Rho1 GTPase (25, 26). Caspase-3 also cleaves and activates gelsolin, an actin-severing and capping protein, thereby altering actin filament turnover (27).

Our previous work has shown that Drice (Drosophila caspase-3 homolog), is necessary for the precise development of MTs, which are primarily associated with excretion and the osmoregulation, similar to human kidneys (28). The MTs, despite expressing active caspase-3, do not undergo apoptosis during metamorphosis, unlike most of the other larval tissues, and are carried into the adult stage. Notably, in caspase-3/Drice–deficient flies, MTs show disrupted actin architecture, characterized by abnormally dense F-actin accumulation and multiple fluid filled cysts, shorter tubule length and functional impairment (28). This phenotype is accompanied by dysregulated Rho1 GTPase expression, suggesting a link between caspase signaling and Rho1-mediated cytoskeletal organization (28). However, the exact role of caspases in the modulation of *Rho1GTPase-*mediated cytoskeletal regulation, as well as, other underlying reasons for the failure to achieve normal morphological characteristics of MTs, remained a question.

In this study, we provide a comprehensive analysis of how two members of the RhoGTPase family, Rho1 GTPase and CDC42 are altered in the absence of Drice and how this disruption affects tubule morphology in MTs of *Drosophila melanogaster*. We show that in the absence of the Drice the downstream effector of Rho1, i.e., Rok expression is reduced, which eventually leads to the trs/cofilin activation and increases actin polymerization. On the other hand, increased expression of CDC42 in the MTs results in enhanced Arp2/3 expression consequently increasing actin branching. In addition, we demonstrate that gelsolin (a key actin-severing protein required for filament clearance) is downregulated in the absence of caspase-3, also resulting in excessive F-actin accumulation and cytoskeletal disorganization. The cumulative effect of these signaling errors results in an imbalance in F-actin to G-actin ratio. In addition to redefining the cytoskeletal changes caused by caspase-3, this work highlights the broader role of caspases in developmental and physiological processes beyond cell death.

## Result 1: Rok downregulation disrupts actin organization in MTs of Drice mutant

In our previous study, we reported that homozygous mutation in Drice, *w^1118^;+/+;Drice*^Δ*2C8*^*/Drice*^Δ*2C8*^, results in abnormal development of the MTs. They exhibit distorted morphology along their length with numerous fluid-filled cysts (28). Drice mutant’s Malpighain tubules develop extensive accumulation of actin filaments and enhanced *Rho1GTPase* expression (Supplementary figure S1) as reported earlier (28). The question that we are now addressing is whether actin dynamics is modulated by the disruption of Rho1GTPase alone, or the other members of the Rho family *viz.,* Rac and CDC42, are also perturbed in absence of the Drice. We first examined the expression of Rok, the immediate downstream effector molecule of Rho1GTPase, by qPCR in the MTs from 3rd instar larvae. Transcript analysis by qPCR revealed a significant reduction in Rok mRNA levels in Drice^Δ2C8^/Drice^Δ2C8^ larvae compared with *Oregon R+* (Fig. 1, A). In order to find whether reduced mRNA levels also resulted in reduced protein levels, immunostaining was performed with anti-Rok antibody. A distinct reduction was visible in the MTs of Drice^Δ2C8^/Drice^Δ2C8^ (Fig. 1, B-b, d) compared with *Oregon R+* (Fig. 1, B-a, c). Rok remains in self-inhibitory state under basal conditions. It is activated either through Rho1 association at the Rho-binding domain or via proteolytic cleavage of the C-terminal cysteine-rich region by caspase-3 (25, 29, 30). Western blot analysis using a anti-Rok antibody further confirmed a significant reduction in Rok protein levels in *Drice*^Δ2C8^/*Drice*^Δ2C8^ larvae (Fig. 1, C). To further understand the role of Rok in the MTs, Rok was downregulated by employing UAS-Gal4 system (31). MT specific Gal4, *c42*, was used to express *UAS-Rok RNAi*. It was observed that in *c42-Gal4>UAS-Rok RNAi* progeny, F-actin filaments appeared to be thicker and the regular arrangement was lost (Fig. 1, D-c, m; yellow arrows) as compared to *Oregon R+* (Fig. 1, D – a & k), but not to the extent of Drice (Fig. 1, D – b, l). Additionally, Rok knockdown *viz., c42-Gal4>UAS-Rok RNAi* also resulted in extra budding and morphological defects in the tubules (Supplementary Fig. S2), indicating a direct role of Rok in tissue morphogenesis. Having observed that knockdown of Rok disrupted actin organization, we overexpressed Rok in the Drice mutant background using MT specific UO-Gal4. As expected in the MTs of *UO-Gal4/+; Drice*Δ*2c8/UAS-Rok,* progeny, a significant reduction of cytosolic F-actin accumulation along with restoration of F-actin organization resembling *Oregon R+* was observed (Fig. 1, D-d, n). The morphology of the tubules also reverted similar to *Oregon R+* (Fig. 1, D-a, k). On the contrary, Rok knockdown in the Drice mutant background (*UO-Gal4/+; Drice*Δ*2c8/UAS-Rok-RNAi*) led to further disruption of F-actin in the MTs (Fig. 1, D-e, o).

**Figure 1:**
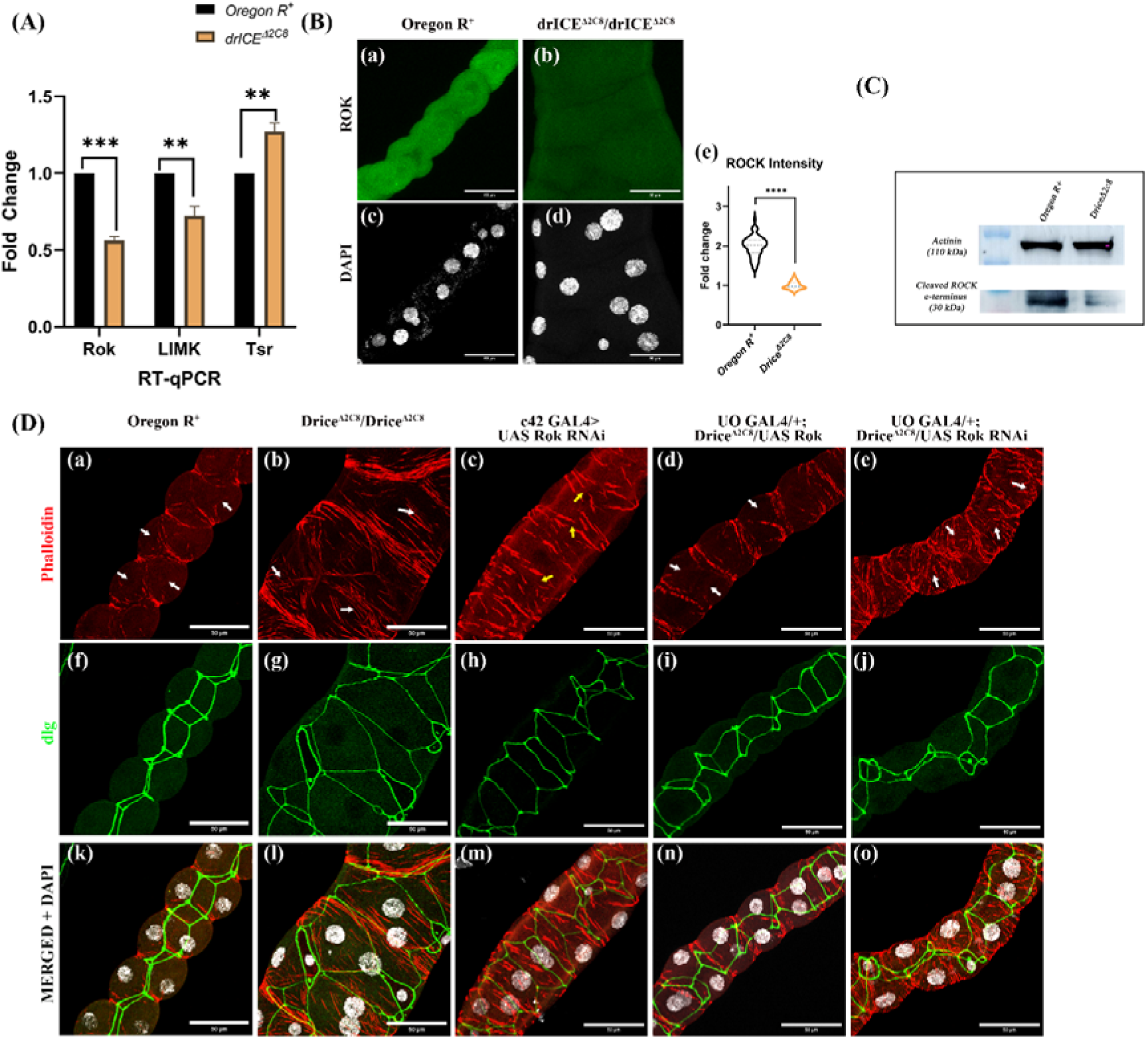
Downregulation of Rok leads to cytosolic actin disruptiion. (A) Transcript levels of *Rok, LIMK*, and *tsr* in third instar larvae. (normalized to *Rp49* as internal control). *Rok* and *LIMK* transcripts are significantly downregulated, whereas *tsr* is significantly upregulated in Drice^Δ2C8^/Drice^Δ2C8^ larvae compared to *Oregon R+.* Unpaired student’s t-test was done to determine the statistical significance. p-value ≤ 0.05 is considered significant, with reference *p < 0.05, ** p <0.01, ***p < 0.001, and ****p < 0.0001. Bar graph is showing Mean ± SEM value. **(B) Rok protein expression in Drice muatnts:** Drice^Δ2C8^/Drice^Δ2C8^ larvae show markedly reduced Rok protein expression (B-b, d) compared to *Oregon R+* (B-a, c). Quantification of fluorescence intensity is shown in histogram (B-e). Magnitude of scale bar is 50 µm. Unpaired student’s t-test was done to determine the statistical significance. p-value ≤ 0.05 is considered significant, with reference *p < 0.05, ** p <0.01, ***p < 0.001, and ****p < 0.0001. Bar graphs are showing Mean ± SEM value. **(C) Western blot analysis of Rok protein.** Caspase-3 mediated C-terminus cleaved Rok (∼30 kDa) is strongly reduced in Drice^Δ2C8^/Drice^Δ2C8^ larvae compared to *Oregon R+*. Actinin (∼110 kDa) was used as loading control due to variability in actin and tubulin levels between samples (34). **(D) F-actin organization (in red) and apico-basal polarity (green) in the MTs:** *Oregon R+* MTs display continuous cortical F-actin fibers with clear cytosolic regions (D-a, f, k; white arrows). In contrast, Drice^Δ2C8^/Drice^Δ2C8^ larvae exhibit dense and disorganized/condensed cytosolic F-actin accumulation with disruption of apico-basal polarity (D-b, g, l; white arrows). Knockdown of Rok *(c42-Gal4>UAS-Rok-RNAi*) results in thick, discontinuous F-actin fibers (D-c, h, m; yellow arrows). Overexpression of Rok in the Drice mutant background (*UO-Gal4/+; Drice*^Δ*2C8*^*/UAS-Rok)* restores F-actin localization and dlg expression (D-d, i, n) similar to *Oregon R+*, whereas Rok knockdown in the Drice mutant background (*UO-Gal4/+; Drice*^Δ*2C8*^*/UAS-Rok-RNAi*) further exacerbates F-actin disorganization (D-e, j, o). Magnitude of scale bar is 50 µm.

Discs large (Dlg) is a junctional protein that marks the basolateral membrane and is essential for maintaining apico–basal polarity of the tubules (32, 33). In Drice mutants, Dlg expression was severely disrupted (Fig. 1, D-g), reflecting disruption in the morphology compared with wild-type (Fig. 1, D-f). Similar to Drice, Dlg disruption was observed in *Rok-RNAi* (Fig. 1, D-h); however, this defect was rescued in *UO-Gal4/+; Drice*Δ*2c8/UAS-Rok* progeny (Fig. 1, D-i). Taken together, these findings suggest that the high cytosolic F-actin accumulation in *Drice* mutant MTs results from an imbalance in the downstream effectors of the Rho1 pathway. These findings suggest potential role for Rok in regulating morphogenesis of the MTs.

## Result 2: *CDC42*, is necessary for appropriate actin organization in the MTs

Given the dysregulation of the Rho1 downstream cascade, we next examined the status of other Rho GTPase family members, namely Rac and CDC42. The interaction between Rho1 and Rac is known to be context dependent, leading to either mutual activation or antagonism depending on the dominant Rho1 target within the cell (35). In contrary, CDC42 has been identified as a key inhibitor of Rho1 activation during cytokinesis in early anaphase (36). qPCR was done to check the status of transcription of Rac and CDC42 in the MTs of 3rd instar larvae. Transcript analysis did not reveal significant change in *Rac* expression, whereas, *CDC42* was significantly upregulated in *Drice*^Δ*2C8*^*/Drice*^Δ*2C8*^ larvae (Fig. 2, A). Based on these observations, we analyzed the role of CDC42. Immunostaining of 3^rd^ instar larval MTs with anti-CDC42 antibody showed significant increase in CDC42 expression in *Drice*^Δ*2C8*^*/Drice*^Δ*2C8*^ larvae (Fig. 2, B- b & d) compared with *Oregon R+* (Fig. 2, B- a & c). Western blot analysis further confirmed the enhanced expression of the CDC42 protein level (Fig. 2, C), suggesting a potential role for *CDC42*, in the development of the MTs.

**Figure 2:**
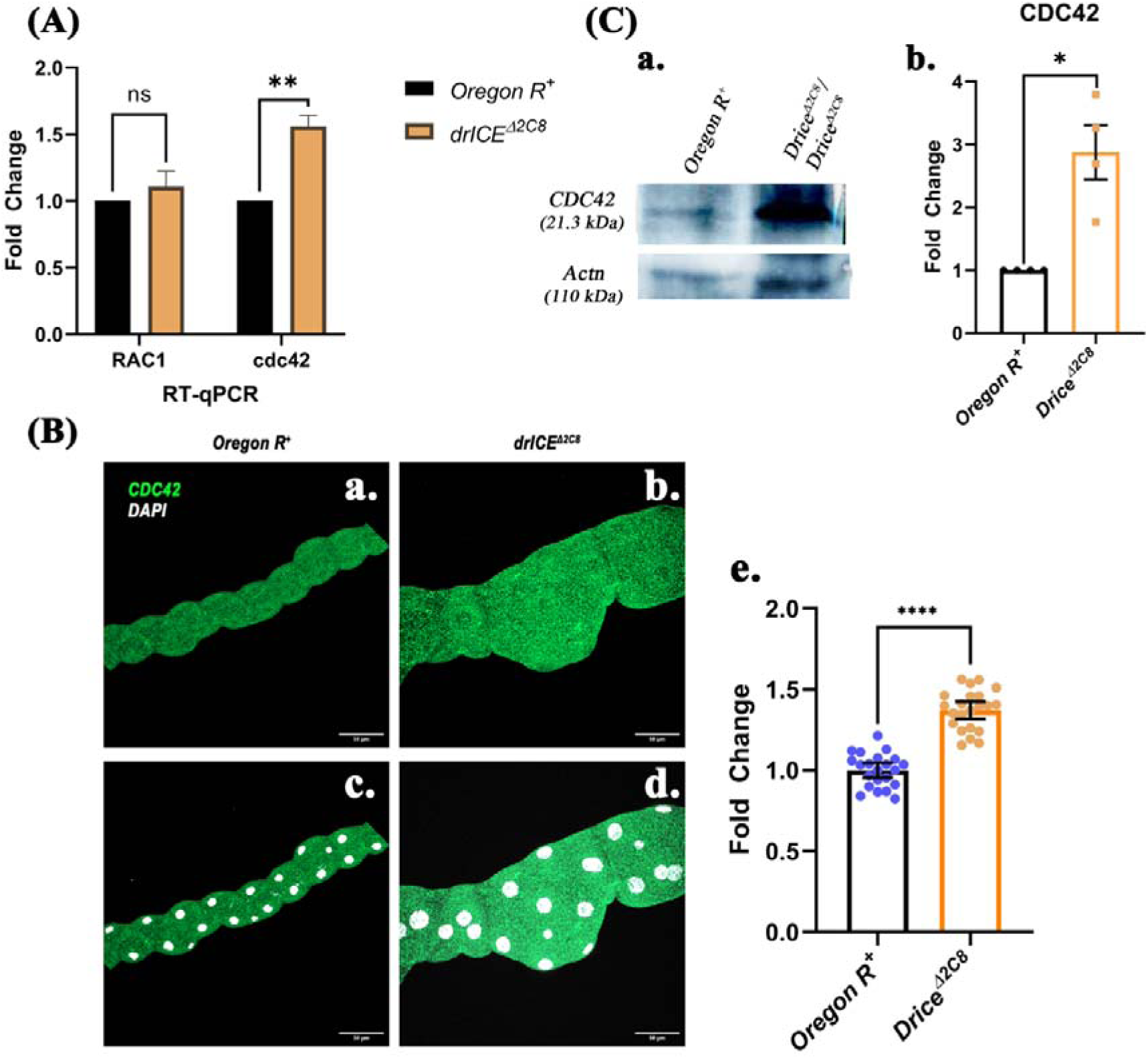
Upregulation of transcript and protein levels of CDC42 in Drice mutants. **(A)** Transcript levels of the *Rac* and *CDC42* in the MTs of 3^rd^ instar Drice^Δ2C8^/Drice^Δ2C8^ larvae when compared to the *Oregon R+* in qRT-PCR (*RP49* was used as internal control). **(B) Elevation of CDC42 in the immunostaining:** CDC42 protein is significantly upregulated in the MTs of wondering 3^rd^ instar larvae of Drice^Δ2C8^/Drice^Δ2C8^ (B-b & d) and in *Oregon R+* (B-a & c) and their quantification (B-e). Scale bar is 50µm. **(C)** Western blot analysis of the CDC42 protein in *Drice*^Δ*2C8*^*/Drice*^Δ*2C8*^ and *Oregon R+* (Actinin was used as internal control antibody). Magnitude of scale bar is 50 µm. Unpaired student’s t-test was done to determine the statistical significance. p-value ≤ 0.05 is considered significant, with reference *p < 0.05, ** p <0.01, ***p < 0.001, and ****p < 0.0001. Bar graphs are showing Mean ± SEM value.

## Result 3: Arp mediated overaccumulation of F-actin at the cell cortex

Having observed upregulation of CDC42, we next investigated the downstream effectors of CDC42 in Drice mutants. CDC42 interacts with WASP, alleviating self-inhibitory state; activated WASP subsequently recruits the Arp2*/*3 complex, promoting nucleation and branching of actin filaments (37–39).

Transcript levels of both Arp2 and Arp3 were significantly upregulated in *Drice*^Δ*2C8*^*/Drice*^Δ*2C8*^ mutants, as shown by qRT-PCR (Fig. 3, B). To assess functional relevance, we downregulated Arp2 by driving *UAS Arp14D-RNAi* with c42-Gal4. In *c2Gal4>UAS-Arp14D-RNAi* progeny the actin filaments appeared to be disrupted, thin and discontinuous in MTs as indicated by arrowheads (Fig. 3, C-c, g, k) compared with *Oregon R+* (Fig. 3, C- a, e, i). Similarly, knockdown of *Arp3, viz., c42- Gal4> UAS-ArpC3A-RNAi* progeny showed actin filaments comparable to that of *Arp2 RNAi* (Fig. 3, D- d, h, l).

**Figure 3:**
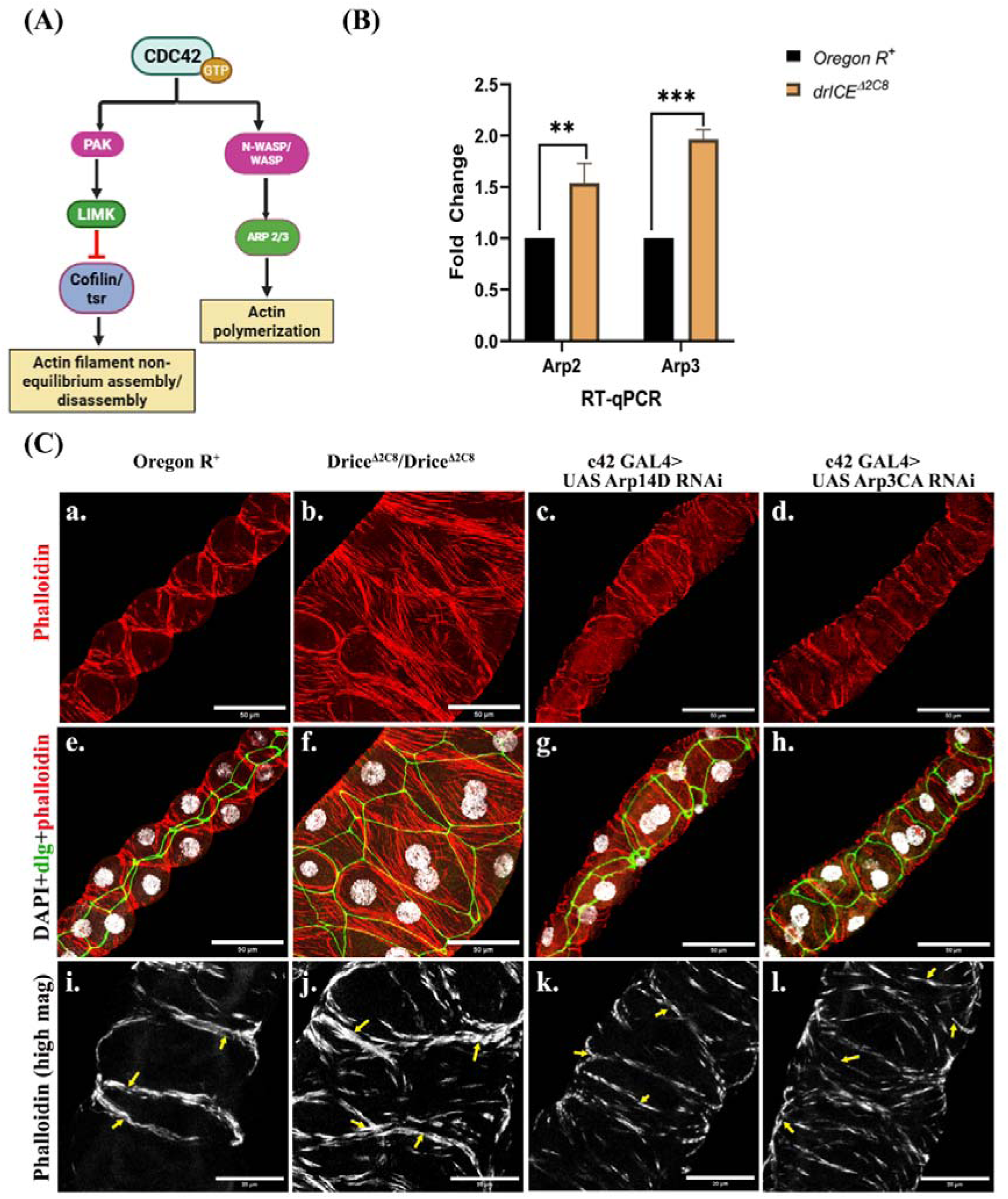
Role of Arp in the actin branching and polymerization. **(A)** Downstream effectors of the CDC42. **(B) Transcript levels of the *Arp2 & Arp3*** in the MTs of 3^rd^ instar Drice^Δ2C8^/Drice^Δ2C8^ larvae when compared to the *Oregon R+* in qRT-PCR (RP49 was used as internal control). p-value ≤ 0.05 was considered significant. Unpaired student’s t-test was done to determine the statistical significance. p-value ≤ 0.05 is considered significant, with reference *p < 0.05, ** p <0.01, ***p < 0.001, and ****p < 0.0001. Bar graphs are showing Mean ± SEM value. **(C) Arp knockdown led to the thinning of the corticle F-actin filaments:** Dense actin fibers at the cell cortex in Drice mutants (C-b, f & j) when compared to the *Oregon R+* actin localization within the MTs (C-a, e & i).

Since Arp2/3 complex creates branching of the actin filaments, it can be hypothesized that their enhanced expression could be responsible for excess networking of actin fibers in Drice (Fig. 3-D, b & j) and reduction in Arp by RNAi (Fig. 3, D- c, d, k, & l) reduces branching. These observations provide strong evidence that Arp2/3 complex has a critical role in the regulation of cortical actin in MTs.

Knockdown of Arp2 (*c42-Gal4>UAS Arp14D RNAi*) (C- c, g & h) led to the incomplete and thinner actin filaments at the cell cortex, also, knockdown of the Arp3 (*c42-GAL4>UAS ArpC3A RNAi*) (C-d, h, & l) led to the thinner actin filaments at the cell cortex in the MTs. Fig a-h is showing 40x 1.0 zoom image; red, green and gray are showing phalloidin, dlg and DAPI respectively (scale bar = 50µm). Fig i-l are showing 63x 1.5 zoom image showing phalloidin staining in gray (scale bar = 20µm). Arrow head is showing the F-actin fibers thickness in different genetic background.

## Result 4: Gelsolin a mediator between Drice and Rho1GTPase, in regulation of the actin dynamics within the MTs

After exploring the roles of *Rho1-GTPase* and *CDC42* in actin dynamics in MTs, a critical question remained: what role does *Caspase-3/Drice* play in coordination with *Rho1* family members in regulating the actin cytoskeleton? To address this, immunoprecipitation-mass spectrometry (IP-MS) was performed from the larval MTs lysate of *Oregon R+* and Drice mutants using an anti-Rho1 antibody to identify the interacting partners of Rho protein in the different genetic backgrounds. A total of 235 interactors in *Oregon R+* and 264 interactors in Drice mutant were identified, interacting with Rho1. Of these 211 interactors were found in both the genotypes, while 24 interactors are unique in *Oregon R+* and 53 in Drice mutants. Several proteins like troponin, flare, cpa, cpb, Arp3, and tropomyosin-I were unique interactors of Rho1 in Drice mutants, whereas, proteins like gelsolin, myosin, moesin, cora, lva etc., were unique interactors identified in *Oregon R+* (Table 1, Supplementary Data). Among these, gelsolin was of particular interest, as it severs and cap F-actin and it is substrate of the Caspas-3 as well.

To assess potential interactions between Rho1 and gelsolin, we performed co-immunoprecipitation (co-IP) experiments. Immunoprecipitation using an anti-Rho1 antibody followed by immunoblotting with an anti-gelsolin antibody revealed minimal to undetectable association between Rho1 and gelsolin in Drice^Δ2C8^/Drice^Δ2C8^ mutants (Fig. 4, A). This result was verified by reversing the experimental conditions—immunoprecipitating with an anti-gelsolin antibody and probing with an anti-Rho1 antibody. Consistent with the initial observation, very low interaction was observed between gelsolin and Rho proteins in Drice mutants when compared with *Oregon R+* (Fig. 4, B).

**Figure 4:**
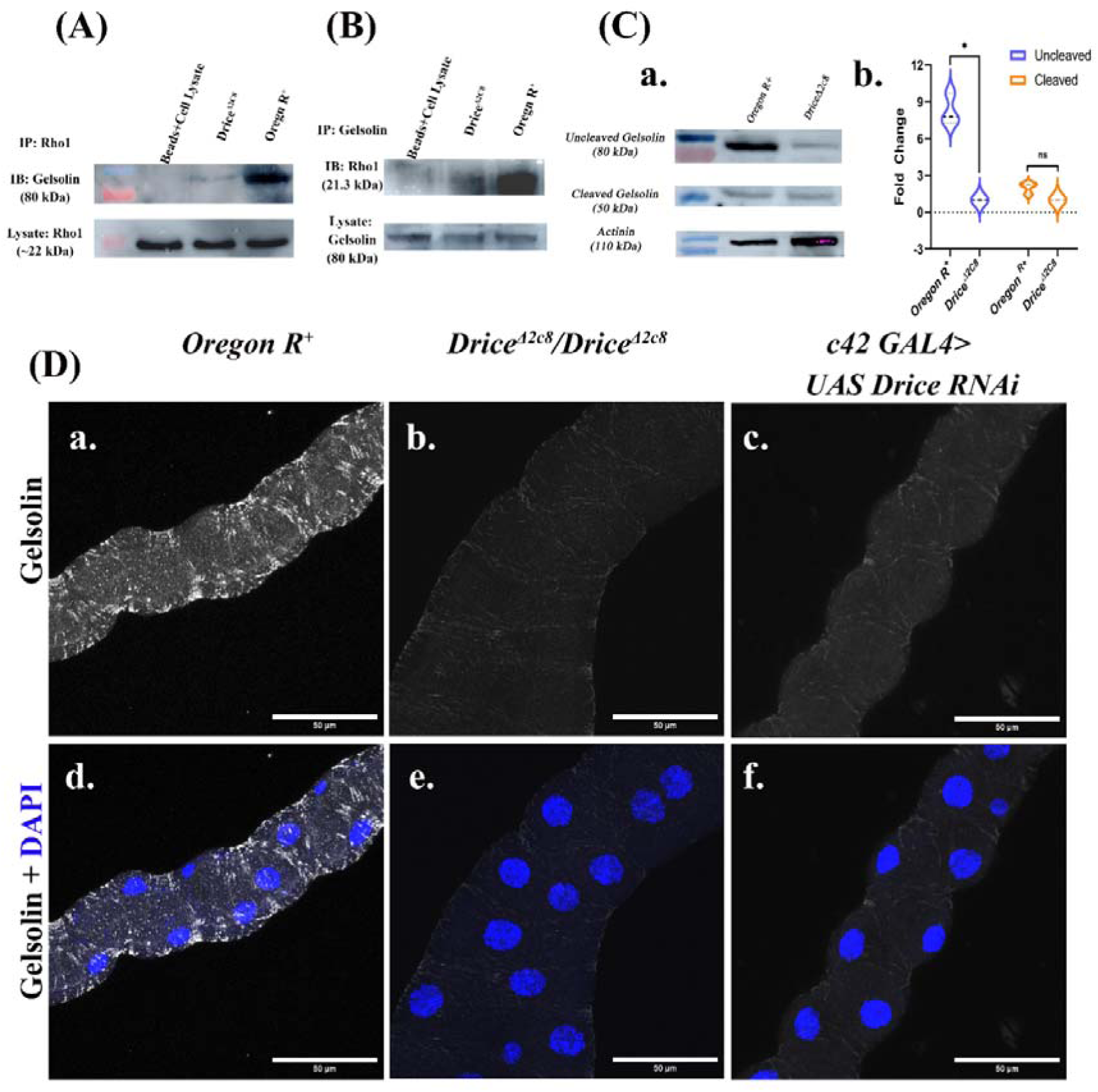
Gelsolin activity in *Oregon R+* and Drice mutants. (A–B) Co-immunoprecipitation (Co-IP) from whole-body lysates of 118–120 h third instar wandering stage larvae using anti- Rho1 or anti-Gelsolin antibodies. (A) Rho1-immunoprecipitated lysates probed with anti-Gelsolin antibody showed no detectable interaction in Drice^Δ2C8^/Drice^Δ2C8^ mutants, whereas the interaction was present in *Oregon R+*. **(B) Gelsolin-immunoprecipitated lysates** from Drice^Δ2C8^/Drice^Δ2C8^ mutants showed no interaction with Rho1. In contrast, both Rho1–Gelsolin and Rok–Gelsolin interactions were present in *Oregon R+*. **(C) Western blot analysis of Gelsolin protein using anti-Gelsolin antibody**. Uncleaved Gelsolin (∼80 kDa) was significantly reduced in Drice^Δ2C8^/Drice^Δ2C8^ mutants compared with *Oregon R+*. A baseline level of cleaved Gelsolin (∼40 kDa) was detected in both genotypes. Actinin (∼110 kDa) was used as loading control. *Unpaired student’s t-test was done to determine the statistical significance. p-value* ≤ *0.05 is considered significant, with reference *p < 0.05, ** p <0.01, ***p < 0.001, and ****p < 0.0001. Bar graphs are showing Mean ± SEM value.* (D) Immunostaining of MTs using anti-Gelsolin antibody (grey). Gelsolin levels were markedly reduced in *Drice*^Δ*2C8*^*/Drice*^Δ*2C8*^ mutants (D-b, e) compared with *Oregon R+* (D-a, d). Knockdown of Gelsolin (*c42-Gal4>UAS-Gelsolin RNAi*) showed a similar reduction, consistent with *Drice* mutants. Magnitude of scale bar is 50 µm.

Gelsolin is an ∼80 kDa actin-severing protein that requires Caspase-3–mediated cleavage for activation (27, 40). Upon cleavage, the N-terminal fragment severs and caps actin filaments at the barbed end, thereby preventing further actin polymerization (41–43). Further, we checked the level of Gelsolin protein in Drice mutants and in *Oregon R+*, strikingly, significant reduction of gelsolin expression in Drice mutants (Fig. 4, D-b, e) was found when compared with *Oregon R+* (Fig. 4, D-a, d). *c42 Gal4> UAS Drice RNAi* MTs exhibited a similar reduction in gelsolin protein expression (Fig. 4, D-c, f) as in Drice mutants, further supporting the requirement of Drice for gelsolin expression. Next, we did western blot to further validate these findings, which revealed a significant reduction in the gelsolin protein in the Drice mutants when compared to *Oregon R+* (fig. 4, C- a).

Together, these findings suggest that Caspase-3/Drice activity is required for gelsolin production and activation, and in its absence, gelsolin fails to interact with Rho1. The dense actin filaments observed in *Drice*^Δ*2C8*^*/Drice*^Δ*2C8*^ mutants may therefore result from a loss of gelsolin-mediated actin severing activity.

## Result 5: Caspase dependent regulation of F-actin:G-actin ratio within the MTs

Transitioning between G-actin to F-actin is critical for cell movement related processes. Having observed severe impairment of several proteins involved in actin assembly/disassembly, we wanted to check if it correlated with F-actin to G-actin ratio. To address this question, we quantified the relative abundance of F- and G- actin under different genetic backgrounds (Fig. 5, B). F-actin was quantified by staining with phalloidin and G-actin was stained using DNaseI dye and the ratio was calculated by quantifying the fluorescence intensity. Reduced expression of G-actin was observed in Drice^Δ2C8^/Drice^Δ2C8^ (Fig. 5, A- c2) and *c42GAL4>UAS-Drice RNAi* (Fig. 5, A-d2) as compared to *Oregon R+* (Fig. 5, A-a2) and driver, c42 alone (Fig. 5, A-b2), along with a similar decrease in the F-actin levels in the Drice mutant background (Fig. 5, A-b1). Further, Arp2 knockdown (Fig. 5, A- f1- f3) and Arp3 knockdown (Fig. 5, A- e1-e3) does not affect either of F-actin or G-actin expression levels, however, they affect the F-actin organization as discussed earlier (result 3). In gelsolin knockdown (Fig. 5, A- g1-g3), however, the G-actin was significantly reduced with relatively no change the expression levels of the F-actin. Interestingly, *UO-Gal4/+;Drice*^Δ*2C8*^*/Uas-Rok* (Fig 5, A- h1-h3), increased G-actin levels significantly (Fig. 5, A- h2), along with improvement in the F-actin organization in the MTs.

**Figure 5:**
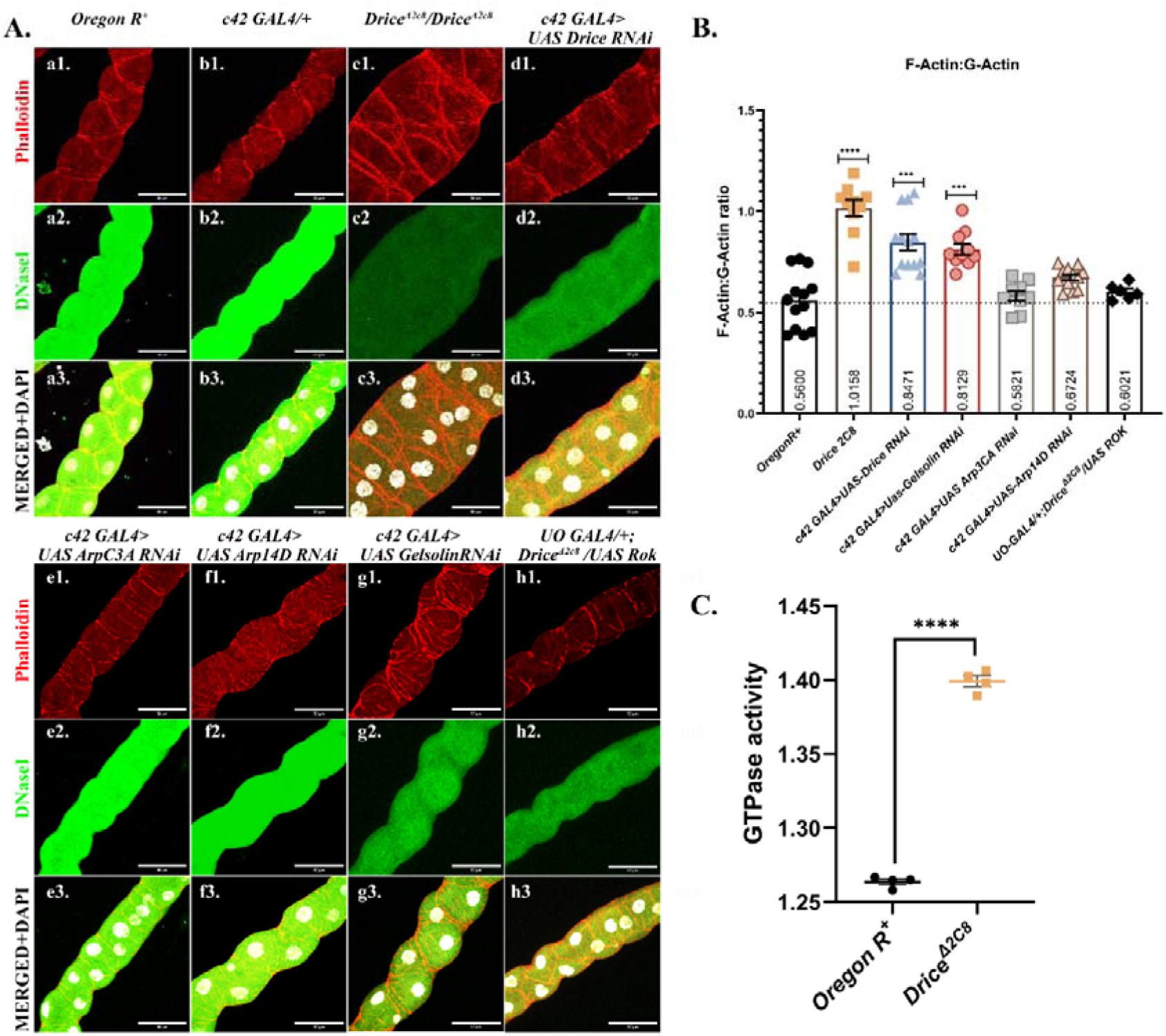
F-actin vs. G-actin dynamics in MTs. (A) Confocal projection images of MTs from wandering third instar larvae (118–120 h) showing F-actin (Phalloidin, red), G-actin (DNaseI, green), and nuclei (DAPI, gray) under different genetic backgrounds. Normal F-actin and G-actin levels in *Oregon R+* (a1-a3) and *c42Gal4/+* driver line (b1-b3); imbalance both in F-actin (organization and level) and G-actin levels in Drice knockdown viz., Drice mutants (c1-c3) and *Drice-RNAi* (d1-d3) can be seen; Arp knockdown viz., *Arp3CA-RNAi* (e1-e3) and *Arp14D-RNAi* (f1-f3) has no effect on the levels of F-actin and G-actin, *Gelsolin RNAi* (g1-g3) reduced G-actin levels significantly with disorganized F-actin, while Rok overexpression in Drice mutant background *viz., UO-Gal4/+;Drice*^Δ*2C8*^*/Uas-Rok* (h1-h3) increases G- actin level with and improvement in f-actin organization similar as in Oregon R+. Magnitude of scale bar is 50µm. (B) Quantification of F-actin and G-actin proportions across genetic backgrounds*: F-/G- actin r*atio is elevated in all *Drice-knockdown* backgrounds (Drice mutants, *Drice RNAi)* and in Gelsolin knockdown (*Gelsolin-RNAi*). Rok overexpression in the Drice mutant background (*UO- Gal4/+; Drice*^Δ*2C8*^*/UAS-Rok*) rescues the ratio to wild-type levels. No significant differences are observed in other genotypes relative to *Oregon R+*. Welch’s one way ANOVA test with Dunnett’s multiple comparison test was done to determine the statistical significance. p-value < 0.05 is considered significant, with reference *p < 0.05, ** p <0.01, ***p < 0.001, and ****p < 0.0001. Bar graphs are showing Mean ± SEM value. (C) Biochemical assay of GTPase activity in MT lysates from wandering third instar larvae (118–120 h). Drice mutants show elevated GTPase activity compared to *Oregon R+*. Unpaired student’s t-test was done to determine the statistical significance. p-value ≤ 0.05 is considered significant, with reference *p < 0.05, ** p <0.01, ***p < 0.001, and ****p < 0.0010. Bar graphs are showing Mean ± SEM value.

Having observed this trend, we looked at F-actin:G-actin proportion into the other genotypes, which affected actin organization. In *Oregon R+*, the F-actin:G-actin ratio in the MTs was ∼0.56 (1:2) (Fig. 5, B), consistent with a balanced state of actin turnover in the absence of active polymerization or depolymerization (44, 45). In contrast, Drice mutants exhibited a marked elevation in the ratio of F- /G-actin ∼1.00 (1:1), directly correlating with enhanced actin fibers. A similar trend was observed in *c42Gal4>UAS-Drice RNAi* (∼0.85). This F-/G- actin ratio imbalance could be due to the absence of Drice. Following the above results, we next sought to look at this ratio in *c42Gal4>UAS-Gelsolin RNAi,* which was ∼0.80 (1:1.25), suggesting enhanced actin polymerization. Similarly, knockdown of *Rok, ArpC3A,* or *Arp14D* did not result in significant deviations from *Oregon R+* F-/G- actin ratio, *viz., c42Gal4>UAS RokRNAi (∼0.58) (1:1.7), c2Gal4>UAS Arp3CA-RNAi (∼0.582) (1:1.7), and c42 Gal4>UAS Arp14D-RNAi (∼0.67) (1:1.5)*; further supporting that these effectors are not the primary determinants of actin remodeling in the MTs.

From the above results, a significant observation was, overexpression of Rok in Drice background (Fig. 5, A, h1-h3) restored F-/G- actin ratio (∼0.60) (1.7) similar to the Oregon R+, suggesting that Rok activation is specifically impaired in the absence of caspase activity. It is known that Rho1 based Rok activation is dependent or/influenced by the cellular GTPase environment, therefore, we biochemically assessed the GTPase activity in Drice mutants. Interestingly, Drice mutants exhibited markedly elevated GTPase activity compared to Oregon R+ (Fig. 5, C). This observation suggests that Rho1 remain in its GDP-bound inactive state and Drice is also absent in Drice mutants, thereby preventing downstream Rok activation. As a consequence, actin hyper-polymerization ensues despite high Rho1 protein abundance.

## Discussion

Microfilaments are one of the major cytoskeletal components that drives tissue architecture. The crucial equilibrium between assembly and dissembly of actin filaments drives the coordinate movement of cells to acquire precise positioning for correct functioning (5, 6). It regulates morphogenetic processes by providing a scaffold that allows cell adhesion, junctional proteins and cell migration, thereby enabling the formation of three-dimensional structures during development and organogenesis (46–48). Several signaling pathways, including small GTPases, PI3K, Hippo, and mTOR, regulate actin dynamics within cells (49, 50). Among these, small GTPase family proteins are key regulators of actin organization, and thus critical determinants of proper cell orientation, movement, morphology, and function (51). Caspases, on the other hand, are traditionally associated with apoptosis and cell death; however, their non-apoptotic roles have recently emerged as an important area of research (28, 52–55).

In the present study we have looked into two of the three small GTPase family, the expression of Rho1 and CDC42 and observed their altered expression in the absence of Drice. Despite high levels of Rho1 in MTs, there was reduced expression of its downstream effector, Rok, which could be because of reduced transcription as observed by qPCR or it could be due to absence of cleavage by Caspase-3, which is known to activate Rok by mediating C-terminus cleavage (25, 26, 56). In the absence of Rok, the inhibition on cofilin/twinstar, which is an essential actin regulatory protein to sever actin filaments accelerating actin assembly dynamics, is removed and it results in increased polymerization of F-actin (57). In Drice mutants, cofilin expression is enhanced (Unpublished data), thus, probably increasing the number of filaments ends from which new actin fibers can polymerize.

The other member of Rho family, CDC42, activates the Arp2/3 complex, which is known to associate with preexisting filaments and leading to the formation of new filaments (58–60). CDC42 promotes polymerization of new actin filaments by initiating actin nucleation or by triggering the uncapping or severing of filaments (61, 62), by activating the Arp2/3 complex through WASP family proteins (63, 64). Enhanced expression of CDC42 in Drice mutants could be responsible for the enhanced expression of WASP (Unpublished data) leading to the activation of Arp2/3 complex. Increased Arp2/3 expression correlates with dense cortical actin fibers (65–67). Conversely knockdown of Arp2/3 led to thinning of cortical actin fibers, indicating that the actin hyper-polymerization observed in Drice mutants is mediated through the Arp2/3 complex.

Rho interaction with gelsolin has not been reported till now, however, this could be a novel observation that in MTs, Rho interacts with gelsolin probably in a tissue specific manner. Gelsolin, being an actin-severing and capping protein regulated either by caspase-mediated cleavage or by calcium signaling (27, 41, 68, 69). In the absence of Caspase-3, Gelsolin remains in its inactive form and fails to sever and cap excess F-actin from the barbed end, thereby exacerbating cytoskeletal imbalances. Thus, Gelsolin could represent a crucial target of Drice that functions in maintaining appropriate F-actin turnover.

The balance between the dynamic states of G-actin and F-actin reflects the physiological status of actin turnover within cells (70). In steady-state conditions, the proportion of G-actin typically exceeds F-actin, with reported F:G ratios ranging from ∼1:1.5 to ∼1:3, indicative of basal equilibrium without active remodelling (44, 45, 71, 72). In contrast, cells undergoing migration display a shift toward higher F-actin abundance relative to G-actin, consistent with active filament assembly (44, 73). The shift in the G- and F-actin pools observed in the absence of Drice could be driving excessive polymerization. F/G-actin ratio further supports these mechanistic insights and is concurrent with an earlier study suggesting this ratio determines the fate of actin polymerization in the biological system (74). Under steady-state conditions without active polymerization, G-actin is higher than F-actin, whereas, high F-actin content represents active polymerization state (44, 45, 71, 72). In accordance with this, high F-actin level (almost same as G-actin) was reported in Drice mutants and Gelsolin knockdown background implicating active actin polymerization in these two. Further quantification revealed, F-actin levels remained largely unaltered across Rok RNAi, Arp3CA-RNAi, Arp14D-RNAi background, however, a notable reduction was observed in Drice mutants and Gelsolin knockdown. G-actin levels were also significantly reduced in Drice RNAi, Gelsolin RNAi conditions, with the most drastic reduction seen in Drice mutants. Importantly, overexpression of Rok in Drice mutants background normalized the F-actin:G-actin ratio by improving the G-actin levels, indicating Rok activity can partially rescue cytoskeletal defects in the absence of caspase activity. This rescue was absent under Rok knockdown, highlighting a unique Rok-dependent regulatory mechanism in Drice mutants. As G-actin acts as the building block for the formation of F-actin (75), and the level of G-actin is very low in Drice and Gelsolin knockdown background, therefore unavailability of G-actin in these genotypes could be the limiting factor for the formation of F-actin. When Rok is overexpressed in the Drice mutant background it increases the G-actin and partially rescued the F-/G- actin ratio in the MTs. Since, Gelsolin is severing and capping F-actin, therefore, in their absence the severing of F-actin in Drice mutants is reduced, further disrupting the F-actin organization. But reason of reduced G-actin cannot be explained, but probably Gelsolin and Drice are involved in maintaining G-actin levels. The failure of triggering of Rho1 downstream effectors despite high Rho1 protein, could be due to the absence of Drice. A plausible explanation is that elevated GTPase activity in the Drice- depleted background traps Rho1 in its GDP-bound inactive form, thereby preventing activation of downstream cascades. However, exact mechanistic regulation of the Rho1 in Caspase depleted background is still elusive and yet to be fully understood.

Compelling data presented in this study suggests that Drice could be fundamental in coordinating Rho family signaling, and activity to organize the actin filaments correctly so that MTs acquire their appropriate morphology. The results highlight Caspase-3/Drice as a central regulator of actin dynamics through its dual control of Rok and Gelsolin, with downstream consequences on Rho1 and CDC42 signaling pathways. The interplay between caspase activity and actin regulators ensures proper balance between F-actin and G-actin pools, which in turn influences MTs morphology. Our study indicates that absence of Drice could be altering function several actin assembly proteins and could significantly affect the convergent extension movement necessary for inferred morphogenesis. These findings broaden the functional repertoire of caspases beyond apoptosis, positioning them as critical modulators of cytoskeletal dynamics. Future studies focusing on the precise regulation of Rho1 activity in caspase-deficient backgrounds will be instrumental in fully understanding the crosstalk between caspase signaling, actin remodelling, and microtubule organization.

## Summary

In *Oregon R+* MTs, Drice activates Rok and gelsolin, while reducing CDC42 levels, maintains the dynamic balance of F- and G-actin. In Drice mutants, Rok activation fails and gelsolin remains inactive, while CDC42–Arp2/3 compensates with excessive cortical actin polymerization. This imbalance leads to hyper-dense filaments, polarity defects, and abnormal tubule morphology.

These findings highlight caspases as non-apoptotic regulators of cytoskeletal remodeling and suggest broader implications in other tissues where actin homeostasis and Rho GTPases are critical. Future work could focus on defining how Drice influences Rho1 nucleotide cycling and whether similar caspase–Rho–actin pathways operate in vertebrate systems.

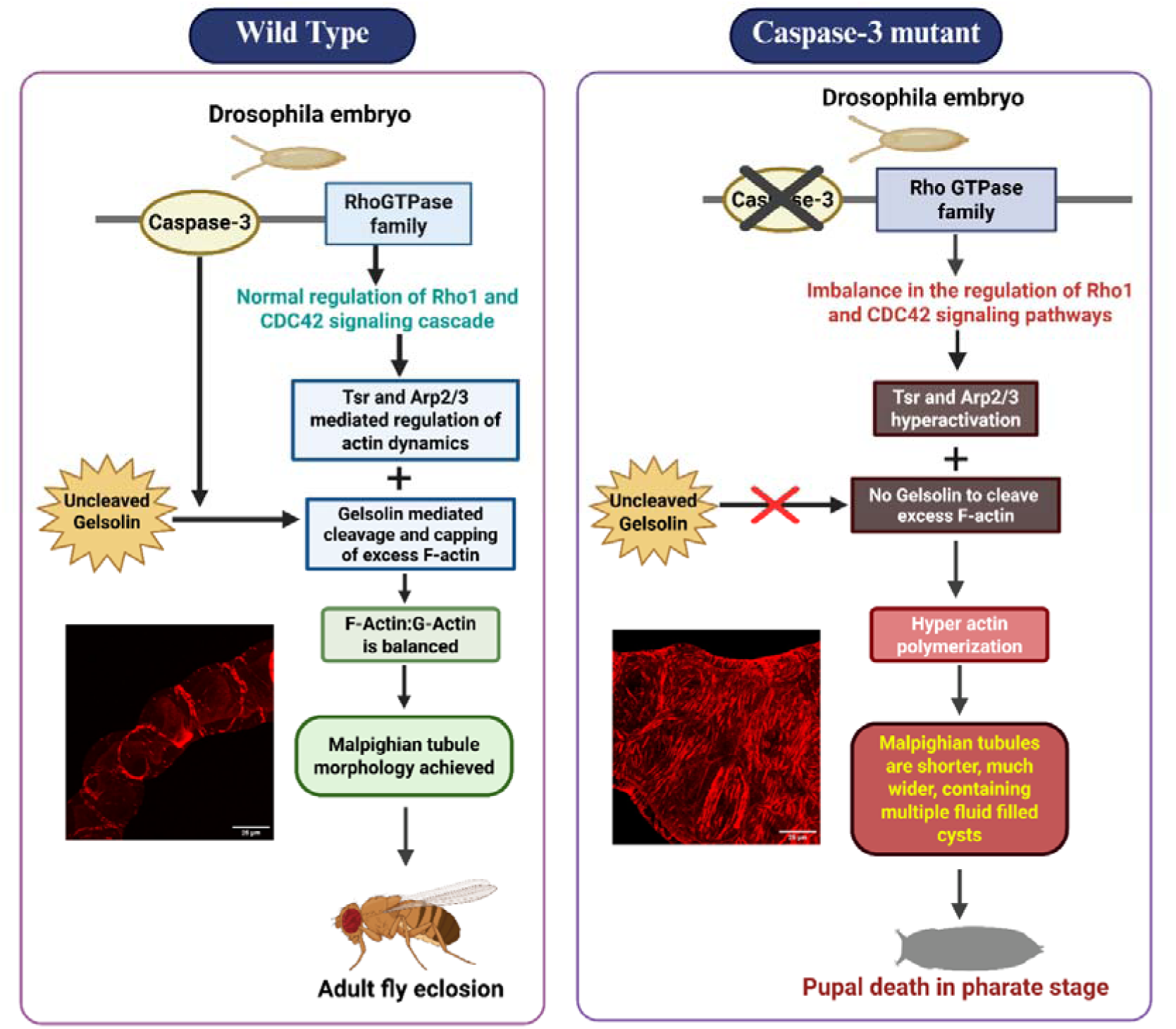

## Material and methods

### 1. Drosophila Stock Husbandry

*Drosophila* stocks were raised on a standard cornmeal fly media containing 0.046% w/v cornmeal, 0.045% w/v sucrose, 0.018% w/v yeast powder, 0.007% w/v agar powder, with additional 0.003% v/v propionic acid and 0.003% w/v hydroxybenzoic acid methyl ester. The stocks were kept in controlled environment with room temperature 25±2°C and experimental flies were reared in BODs, genetic crosses based on Gal4/UAS system were maintained at 29°C under humidity controlled 12-hour light/12-hour dark cycle. The Drosophila used for experiments are: *Oregon R+, Drice2c8, UAS-DriceRNAi (kind gift from Dr. Masayuki Miura lab), UAS RhoGEF2 (BL-9387), UAS Rok 2B1(BL-6670), UAS-Rok RNAi (BL-28797), UAS-mDia RNAi (BL-35479), UAS-Arp14d RNAi (BL-27705), UAS-Arp3CA RNAi (4560R-3), UAS-Gelsolin RNAi (BL31205), c42-Gal4, UO-Gal4 (kind gift from Dr. J.A.T. Dow*.

The MTs of 118-120 hrs old wondering third instar larvae was taken for most of the experiments. Experiments were not sex specific; post eclosion two days old flies were used for genetic cross setup. Min 20 larvae were selected for each experiment and all the experiments were repeated min three times.

### 2. Immunostaining

The MTs were dissected from the wondering third instar larvae in 1xPBS (phosphate buffer saline). Immediately after dissection tissues were transferred in 4% PFA (para formaldehyde) fixative for 45 minutes at room temperature. After fixation tissue were washed in 0.5% PBST (wash buffer) (0.5% Triton X-100 in 1xPBS). Following fixation tissues were incubated in blocking solution (0.1% Triton X-100, 0.1% BSA, 10% FBS, 0.1% deoxycholate, 0.02% thiomersal) for 2 hrs at room temperature and then incubated in primary antibody overnight. Sample were washed thrice in wash buffer afterwards, followed by incubation in blocking solution again for 2 hrs at RT and incubated in the secondary antibody for 2hrs at RT. After incubation in secondary antibody tissues were washed again in 0.5% PBST and counterstained using DAPI (4′,6-Diamidino-2-Phenylindole, Dihydrochloride, Thermo-Fisher Scientific, Cat# D1306) (1 μg/mL) to visualize the nuclei of tissues. Samples were then washed three times and finally mounted in mounting media (DABCO) on glass slide.

Blocking solution was used as dilutant for the antibodies. **The following primary antibodies were used:**

anti-Rho1 antibody raised in mouse (1:20, p1D9, DSHB),
anti-Rho1 antibody raised in rabbit (1:800, 10749-1-AP, Protein tech),
anti-Rho1 antibody raised in mouse (1:300, SC0418, Santa Cruze),
anti-CDC42 antibody raised in rabbit (1:100, 10155-1-AP, Protein tech),
anti-Rok antibody raised in rabbit (1:200, Gly1114, CST),
anti-Gelsolin antibody raised in rabbit (1:200, D9W8Y, CST),
anti-dlg antibody raised in mouse (1:20, 4F3, DSHB).

**The secondary antibodies used for the immunohistochemistry are as follows:**

donkey anti-mouse AF 555 (A31570),
goat anti-mouse AF 647 (A21050),
donkey anti-rabbit AF 555 (A31572) and
goat anti-rabbit AF 647 (A32733) from Invitrogen.

### 3. Phalloidin staining

MTs from 118-120 hrs old wondering stage larvae were dissected in chilled 1xPBS and fixed in 4% PFA for 45 minutes at RT. Tissues were washed thrice in wash buffer (0.5% PBST in 1xPBS) and incubated in phalloidin atto 550 (Sigma) for 2 hrs. Tissues were washed again thrice in washing buffer and counterstained with DAPI for 20 minutes at RT. Tissues were washed and mounted in mounting media (DABCO) on a glass slide and visualized under the microscope.

### 4. DNaseI Staining

MTs from 118-120 hrs old wondering stage larvae were dissected in chilled 1xPBS and fixed in 4% PFA for 45 minutes at RT. Tissues were washed thrice in wash buffer (0.5% PBST in 1xPBS) and incubated in DNAseI Alexa Fluor 488 conjugated dye (D12371, Sigma) for 2 hrs. Tissues were washed again thrice in washing buffer and counterstained with DAPI for 20 minutes at RT. Tissues were washed and mounted in mounting media (DABCO) on a glass slide and visualized under the microscope.

### 5. Western blotting

Third instar Drosophila larvae, aged 119–120 hours post-hatching, were dissected and MTs were collected for protein extraction. MTs were isolated in chilled 1X PBS, then transferred to cold RIPA buffer containing 100 mM Tris-Cl (pH 6.8), 100 mM NaCl, NP-40, EDTA, NaF, NaOLVL, 20% glycerol, a protease inhibitor cocktail, and 2 mM PMSF. Next, the tissues were homogenized and centrifuged at 12,000 RPM for 20 minutes at 4°C, and the resulting supernatant was collected for its protein content. Protein was quantified using Bradford’s assay, and 30 µg of protein was prepared with sample buffer (100 mM Tris-Cl pH 6.8, 4% SDS, 0.2% bromophenol blue, 20% glycerol, 100 mM DTT, and 2 mM PMSF) in a 1:1 ratio. The samples were boiled for 5 minutes and subjected to SDS-PAGE. Proteins were electro transferred onto a PVDF membrane at 100 V for 1.5 hours at 4°C using a wet transfer apparatus. Following transfer, membranes were blocked in 4% BSA dissolved in 0.1% TBST for 2 hours, then incubated overnight at 4°C with the primary antibody. Membranes were washed in 0.1% TBST at RT for 15 minute each washing. They were then incubated with HRP-conjugated secondary antibody for 2 hours at room temperature, followed by another three TBST washes. Signal detection was performed using a ChemiDoc system (Amersion AI680) with ECL substrate (Bio-Rad). All antibodies were diluted in 3% blocking buffer. **The following primary antibodies were used:**

anti-Rho1 antibody raised in rabbit (1:1500, 10749-1-AP, Protein tech),
anti-Rho1 antibody raised in mouse (1:1000, SC-0418, Santa Cruze),
anti-CDC42 antibody raised in rabbit (1:1000, 10155-1-AP, Protein tech),
anti-Rok antibody raised in rabbit (1:1000, Gly1114, CST),
anti-Gelsolin antibody raised in rabbit (1:1000, D9W8Y, CST),
anti-actinin antibody raised in rabbit (1:1000, 2G3-3D7, DSHB).

**The secondary antibodies used for western blotting are as follows**:

HRP tagged anti-mouse IgG (1:1500, Invitrogen)

HRP tagged anti-mouse IgG (1:1500, Invitrogen)

### 6. Immunoprecipitation and immunoblotting

Third instar larvae of Drosophila, 119–120 hours after hatching, were dissected to obtain MTs for protein extraction. The MTs were washed in chilled 1X PBS and then moved to cold RIPA buffer containing 100 mM Tris-Cl (pH 6.8), 100 mM NaCl, NP-40, EDTA, NaF, NaOLVL, 20% glycerol, a protease inhibitor cocktail, and 2 mM PMSF. The tissues were subsequently homogenized and centrifuged at 12,000 RPM for 20 minutes at 4°C, after which the supernatant containing protein was recovered and subjected to IP. For IP 20µl of the Protein G Plus-Agarose Suspension beads (IP04, Miilipore) was taken and washed thrice in 1xPBS at 4LC. After washing beads were incubated with desired primary antibody (5-10 µg in 500 µl 1xPBS) for one hour at RT. Following the incubation, antibody was restored and bead-antibody was washed thrice in 1xPBS at 4^D^C. Antibody incubated beads were then added in 1-2 mg protein and left for 2 hrs at 4^D^C. These beads were again washed 3-5 time in 1xPBS and subjected for immunoblotting or mass-spectroscopy based upon the desired outcome. **The following antibodies were used for IP:** rabbit anti-Rho1 antibody (1:1500, 10749-1-AP, Protein tech), mouse anti-Rho1 antibody (1:1000, SC0418, Santa Cruze), rabbit anti-Gelsolin antibody (1:1000, D9W8Y, CST).

### 7. GTPase assay

GTPase activity was assayed using MAK113 kit (Sigma) as described by the manufacturer. Protein samples from wondering third instar larvae were extracted in Ringer solution (phosphate free buffer, Millipore) and quantified using Bradford assay. Before GTPase assay standard curved was drawn using known phosphate standard and their incubation with reagent provided (hypersensitive malachite green). For biological samples, protein samples were taken and assay buffer was added to make overall volume 10 µl and left for 30 minutes incubation at RT. Following this, reagent (200 µl, malachite green) was added and incubated again for 30 minutes and absorbance was recorded at 620 nm using UV-Vis spectrophotometer. Unknown value was plotted against standard curve and thus determining the GTPase activity in the sample.

### 8. RNA Isolation and cDNA preparation

MTs from 118±1 hrs old post eclosion larvae were collected and immediately processed in TRI reagent (T9424, Sigma Aldrich) and stored at −80 °C until extraction. Total RNA was isolated using the TRIzol method, following the manufacturer’s protocol (Sigma-Aldrich, India), including DNase I (Thermo Fisher Scientific, Cat# 89836) digestion step to remove genomic DNA. RNA concentration and purity were measured spectrophotometrically (A260/A280 and A260/A230 ratios), and integrity was assessed by electrophoresis. First-strand cDNA was synthesized from 200–500 ng total RNA in 20 µL reactions using a reverse transcriptase with a mixture of random hexamers and oligo(dT) primers, following the enzyme supplier’s protocol. Reactions were incubated at 25 °C for 10 min (primer annealing), 42–50 °C for 30–60 min (extension), and heat-inactivated at 70 °C for 10 min. cDNA was diluted 1:5–1:20 in nuclease-free water for qPCR; identical treatment was applied to −RT control reactions.

### 9. qRT-PCR & primer used

Quantitative PCR was performed using a commercial 2× qPCR master mix (SYBR Green) in optical 96-well plates. Reaction composition for a 10 µL SYBR Green (SYBR Green, Genetix, Cat# PKG025-A) reaction was: 1× master mix, 200–400 nM each primer, and 1–2 µL diluted cDNA (final volume 10 µL). Plates were sealed, briefly centrifuged, and run on a real-time PCR instrument. Typical cycling conditions were: 95 °C for 2–3 min (activation), followed by 40 cycles of 95 °C for 10 s and 60 °C for 20–30 s with fluorescence acquisition at the annealing/extension step. A melt-curve was acquired for SYBR assays (65–95 °C, incremental steps). No-template controls (NTC) and −RT controls were included for every primer pair. Each biological condition included at least three independent biological replicates. Technical replicates (duplicate or triplicate wells) were measured for each sample-gene combination. Reference gene stability was assessed across all samples and conditions; RP49 and GAPDH were use used as reference genes and their geometric mean used for normalization. Wells with atypical amplification curves, high replicate variance (>0.5–0.7 Ct), or NTC amplification were inspected and excluded when justified.

**Primers used: following primers were used for qRT-PCR**

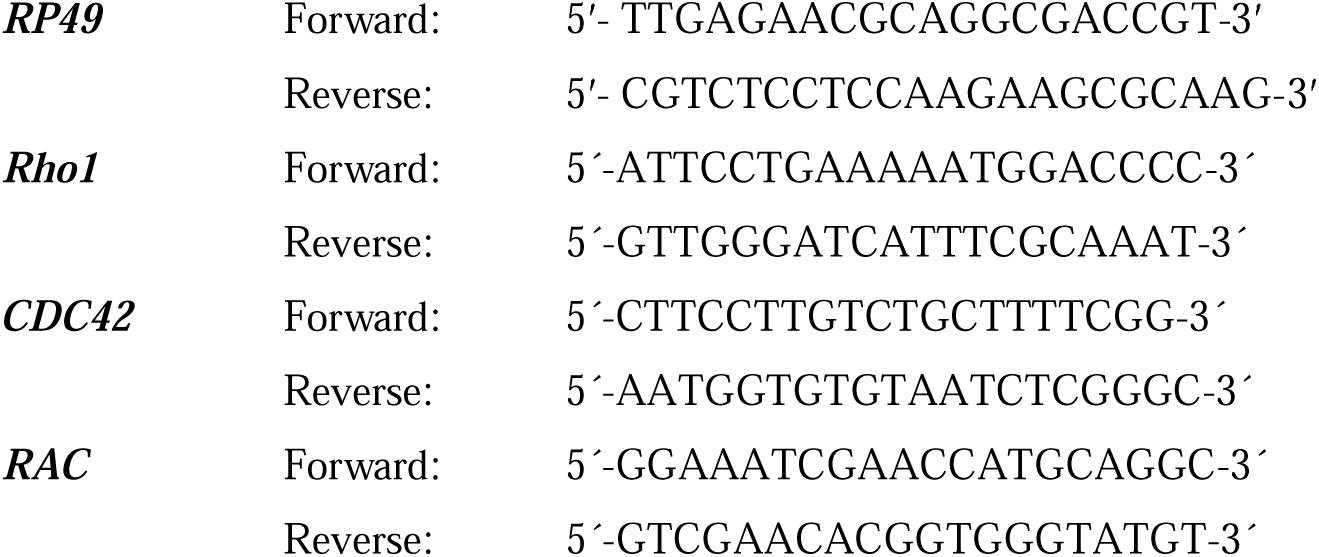
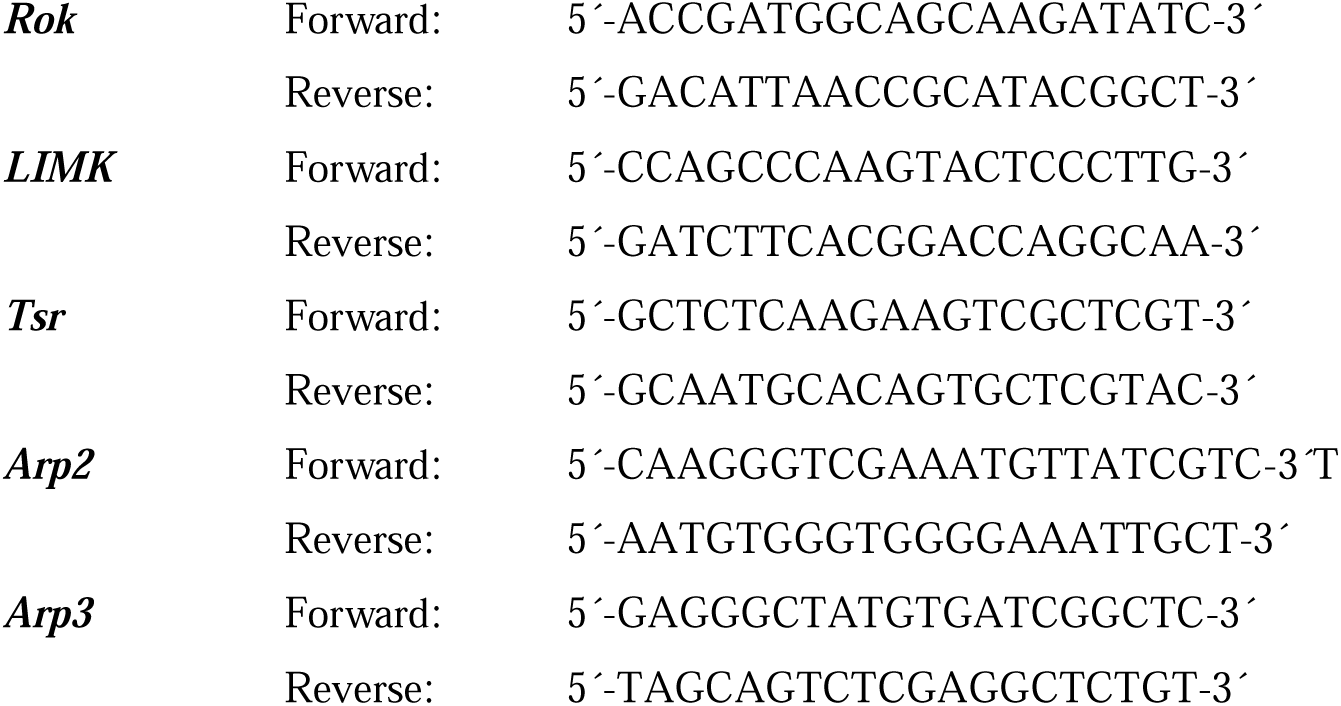

## 10. Microscopy and Image analysis

All the images were scanned using Zeiss LSM-900 confocal microscope using Zen software (version 3.4). For imaging 20x objective and 40x objectives were used on 1x zoom and 63x objective on 1.5x zoom were used, with 2.5 µm optical section interval. Optimum setting was used for image acquisition on different days. Scanned images were processed using ImageJ software (NIH, USA) (available at ImageJ.nih.gov/ij) and Adobe Photoshop 2021 (version 22.4.2) was used to create image panels.

## 11. Statistical Analysis

All the experiments were repeated minimum three times or more and each time a minimum of 10 tissues were taken for the experiment. All the statistical analysis was performed using GraphPad Prism 9. Unpaired two-tailed Student’s t-tests, Welch’s one way ANOVA test was performed in appropriate experiments. p-value ≤ 0.05 was taken as significant difference and degree was significance was as follow: p-values ≤0.05, ≤0.01, ≤0.001, and ≤0.0001 are signifying*, **, ***, and ****, respectively; with p-value > 0.05 representing non-significant statistical difference.

## Acknowledgements

We acknowledge the Department of Zoology, Institute of Science, Banaras Hindu University, Varanasi, for providing basic facility and infrastructure. We thank Dr. J. A. T. Dow, Institute of Biomedical Science, University of Glasgow, U.K. - c42-Gal4; Dr. M. Miura, Department of Genetics, University of Tokyo; for DriceΔ2c8 stocks; and Dr. Pralay Majumder, Presidency University, Kolkata, India for Rok-RNAi, Gelsolin-RNAi, Arp3CA-RNAi, Arp14D- RNAi, UAS-Rok and UAS-Rho GEF2fly stocks. We acknowledge the Interdisciplinary School of Life Sciences, BHU, for Confocal microscopy facility and the Department of Zoology, BHU for the SEM facility. We acknowledge central discovery centre (CDC) SATHI BHU, for IP-MS facility.

---

**Supplementary Figure S1.**
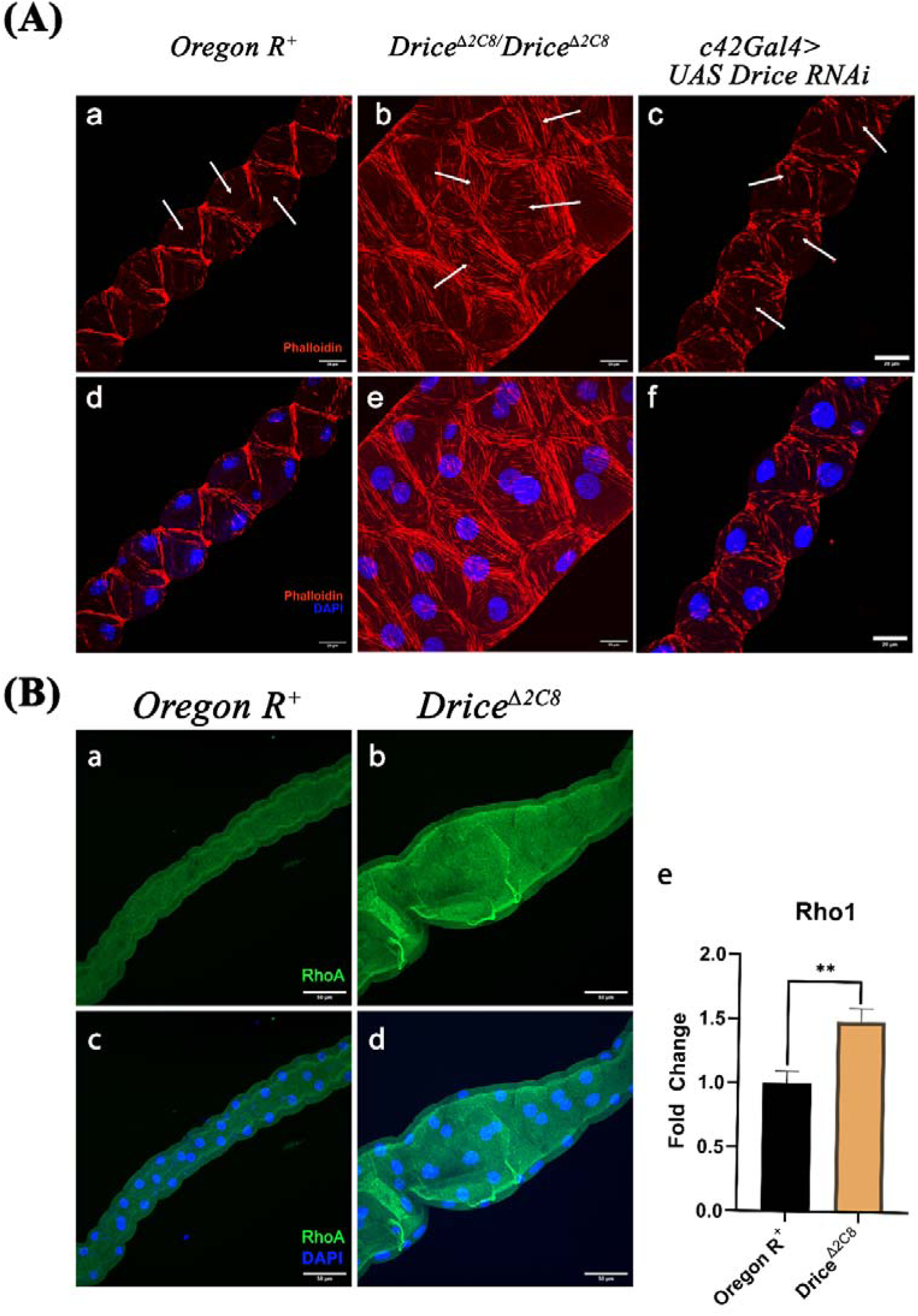
Status of F-actin (red) and Rho1 (green) in Oregon. **R**D **and Drice backgrounds**. In Drice mutants (A-b, e) and *Drice RNAi* (A-c, f), F-actin shows marked disorganization with abundant cytosolic actin fibers compared to the control. Furthermore, Rho1 protein expression is significantly elevated in Drice mutants (B-b, d, e).

**Supplementary Figure S2:**
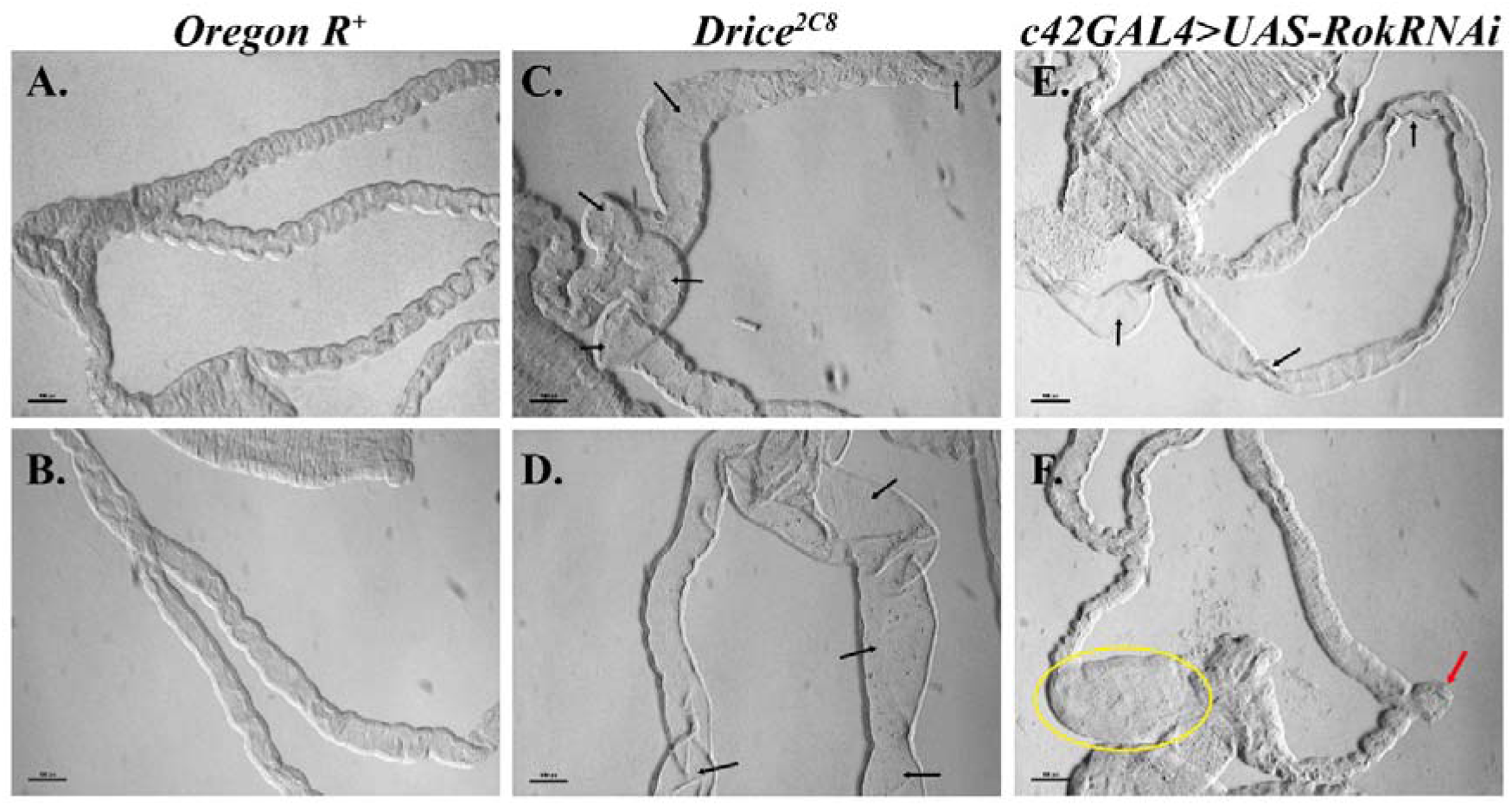
Morphological defects in Drice mutants (C, D) and Rok RNAi background (E, F). Black arrows indicate swollen tubules; the yellow circle highlights the balloon-like swollen ureter region; and the red arrow marks the incomplete tubule arm observed in the Rok RNAi background. Scale bar: 100 µm.

**Supplementary Figure S3.**
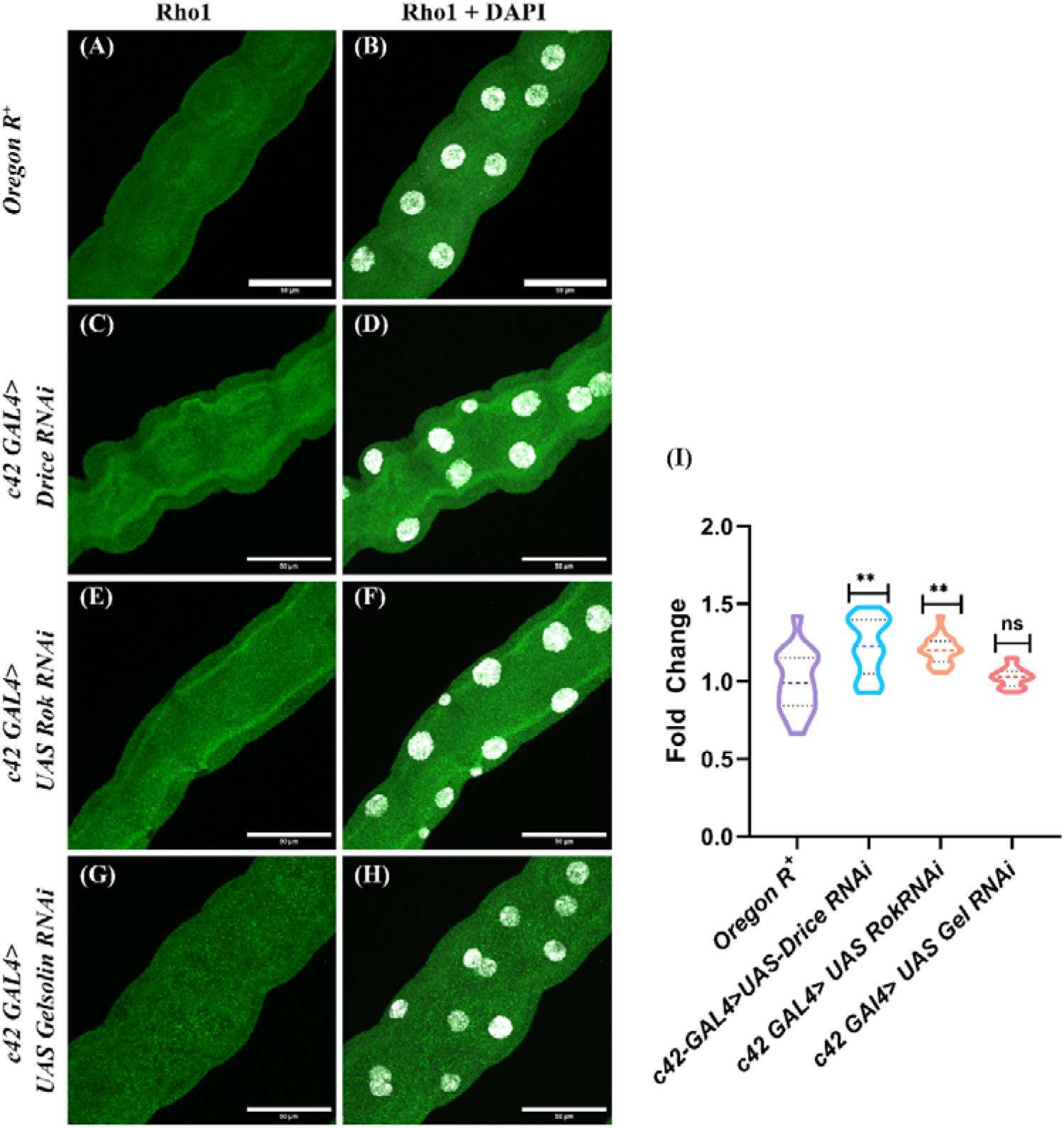
Rho1 protein expression is significantly upregulated in *Drice RNAi* and *Rok RNAi* backgrounds, whereas *Gelsolin RNAi* shows no significant change in Rho1 expression compared with Oregon RL.

**Supplementary Figure S4.**
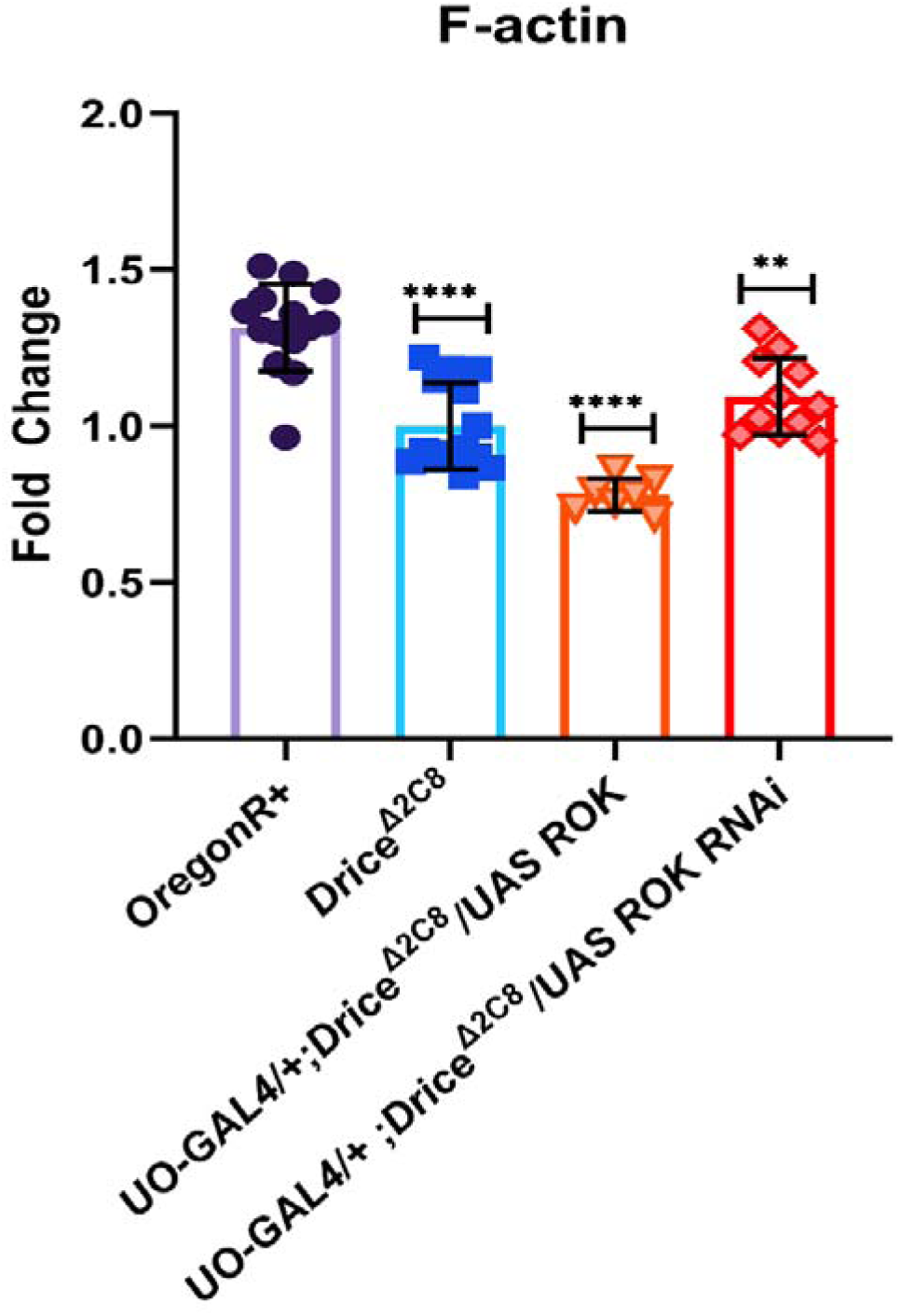
Overexpression of UAS-Rok in the Drice background (*UO-Gal4/+; Drice*Δ*2C8/UAS-Rok*) reduces F-actin levels. In contrast, Rok knockdown in the same genetic background (*UO-Gal4/+; Drice*Δ*2C8/UAS-Rok RNAi*) elevates F-actin levels, although not significantly, possibly due to limited G-actin availability in the Drice mutant background. These results suggest a role for Rok in actin polarization.

**Supplementary Figure S5.**
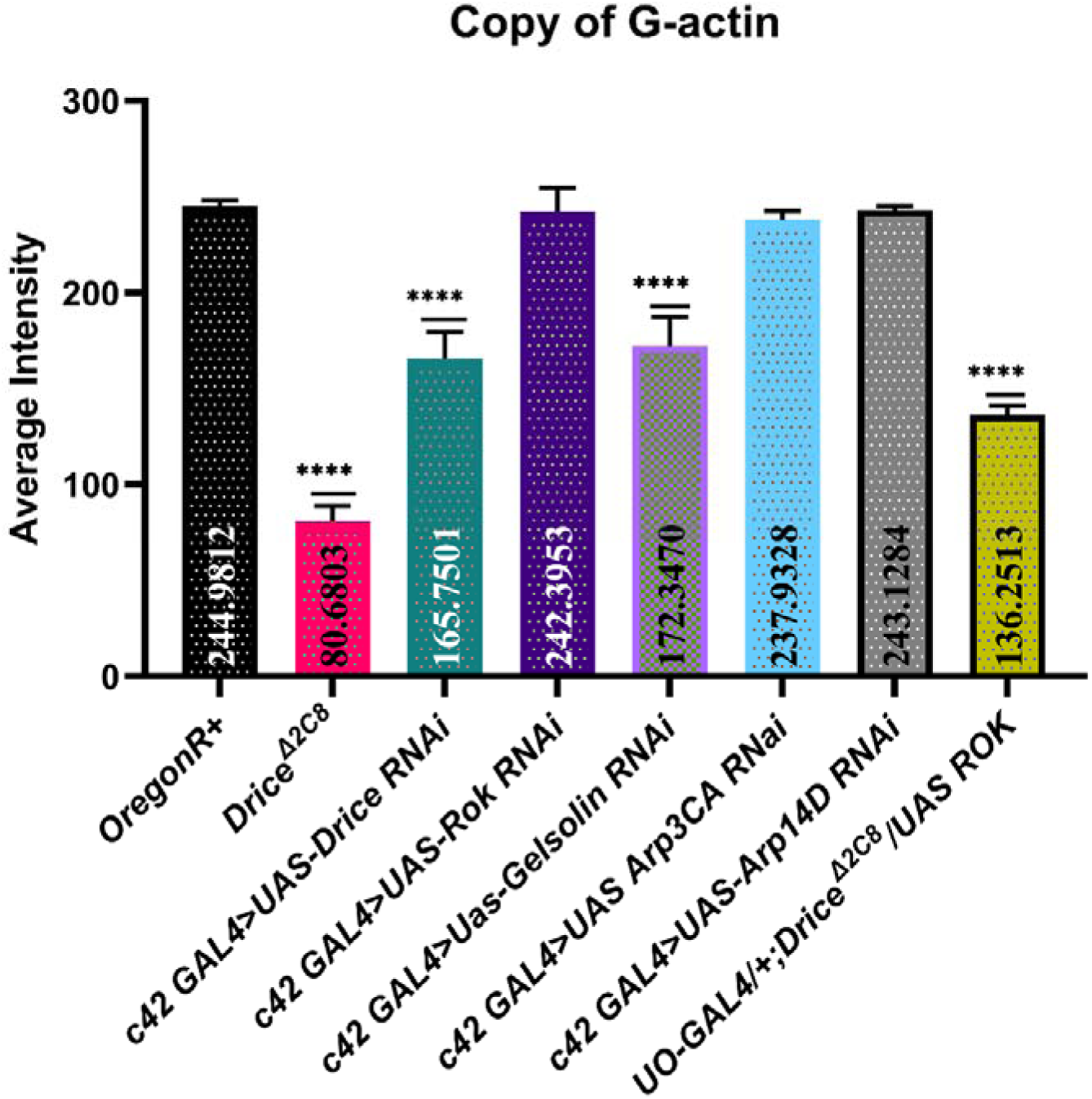
Drice and Gelsolin knockdown significantly reduced G-actin levels, with the most pronounced decrease observed in the Drice mutant background. In contrast, *Rok, Arp2*, and *Arp3* had no significant effect on G-actin levels. Notably, Rok overexpression in the Drice background (*UO-Gal4/+; Drice*Δ*2C8/UAS-Rok*) increased G-actin levels, potentially due to inhibition of actin polymerization via Twinstar/Cofilin.

